# Hydrogen production in the presence of oxygen by *Escherichia coli* K-12

**DOI:** 10.1101/2022.01.11.475878

**Authors:** George D. Metcalfe, Frank Sargent, Michael Hippler

## Abstract

*Escherichia coli* is a facultative anaerobe that can grow in a variety of environmental conditions. In the complete absence of O_2_, *E. coli* can perform a mixed-acid fermentation that contains within it an elaborate metabolism of formic acid. In this study, we use cavity-enhanced Raman spectroscopy (CERS), FTIR, liquid Raman spectroscopy, isotopic labelling, and molecular genetics to make advances in the understanding of bacterial formate and H_2_ metabolism. It is shown that, under anaerobic (anoxic) conditions, formic acid is generated endogenously, excreted briefly from the cell, and then taken up again to be disproportionated to H_2_ and CO_2_ by formate hydrogenlyase (FHL-1). However, exogenously added D-labelled formate behaves quite differently from the endogenous formate and is taken up immediately, independently, and possibly by a different mechanism, by the cell and converted to H_2_ and CO_2_. Our data support an anion-proton symport model for formic acid transport. In addition, when *E. coli* was grown in a micro-aerobic (micro-oxic) environment it was possible to analyse aspects of formate and O_2_ respiration occurring alongside anaerobic metabolism. While cells growing under micro-aerobic conditions generated endogenous formic acid, no H_2_ was produced. However, addition of exogenous formate at the outset of cell growth did induce FHL-1 biosynthesis and resulted in formate-dependent H_2_ production in the presence of O_2_.

## INTRODUCTION

*Escherichia coli* (*E. coli*) is a **γ**-proteobacterium and facultative anaerobe [1]. The bacterium is commonly found as a gut commensal in many animals (which is an anaerobic environment) [2], but is also found in estuaries (where micro-aerobic environments can be identified) [3], and in addition is associated with plant tissues and other aerobic environments [4]. As a result, *E. coli* can adapt, primarily through regulating gene expression, to changing environmental conditions [5].

The preferred mode of energy metabolism for *E. coli* is aerobic respiration. To do this, short, quinone-dependent, respiratory electron transport chains are assembled [6]. The bacterium has no cytochrome *bc*_1_ complex or cytochrome *c* oxidase, but instead calls upon three quinol oxidases to carry out aerobic respiration: cytochrome *bo*_*3*_ oxidase; cytochrome *bd* oxidase I; and cytochrome *bd* oxidase II. Each of these enzymes has a different affinity for O_2_, with cytochrome *bo*_*3*_ oxidase having the lowest affinity for O_2_ and cytochrome *bd* oxidase II having the highest, thus allowing the bacterium to compete under a full range of O_2_ tensions [7].

A prominent electron donor in aerobic respiration is NADH, however, other possible respiratory electron donors in play under this condition are formate, glycerol-3-phosphate, pyruvate and succinate [6]. In order to use formate (HCOO^-^) as a respiratory electron donor, *E. coli* produces two respiratory formate dehydrogenases: FDH-N and FDH-O [8]. These enzymes oxidise formate at the periplasmic side of the cytoplasmic membrane and pass electrons to the quinone pool thus generating a transmembrane proton gradient by a redox loop mechanism [9]. The FDH-N isoenzyme is encoded by the *fdnGHI* operon and is maximally expressed anaerobically in the presence of exogenous nitrate, while the FDH-O enzyme shares sequence identity with FDH-N, but the genes encoding this isoenzyme are constitutively expressed [8]. Under aerobic (oxic) conditions, FDH-O is thought to couple formate oxidation to the reduction of O_2_ to water by cytochrome *bd* oxidase I [10].

Under anaerobic (anoxic) conditions, and in the complete absence of any other exogenous respiratory electron acceptors, *E. coli* will carry out a ‘mixed-acid’ fermentation of glucose [11]. Here, *E. coli* will generate ATP by substrate-level phosphorylation and excrete ethanol, together with a number of different organic acids, as waste products. These include acetic acid, lactic acid and formic acid. Under fermentative conditions, the *E. coli* TCA cycle is no longer a cycle. Instead, the reactions are split into a reductive arm, terminating with a low level of endogenous fumarate reductase activity, and an oxidative arm, terminating with isocitrate dehydrogenase [12]. Acetyl-CoA is fed into the oxidative arm of the TCA cycle *via* the action of the anaerobic pyruvate formate lyase (PFL) enzyme, which is fully activated in the absence of O_2_. PFL combines pyruvate with coenzyme A to generate acetyl-CoA and a formate anion [13]. Interestingly, the PFL enzyme is physically associated with the formate channel FocA at the cytoplasmic face of the plasma membrane [14]. This means that the formate so-generated is most-likely constantly excreted from the cell *via* FocA in the early stages of fermentation. In a closed batch fermentation environment, the formate (p*K*_a_ 3.7) will accumulate outside the cell together with acetic acid and other products of mixed-acid fermentation, where the pH is thought to tend towards acidic as a result. At around pH 6.8, the FocA channel switches from a facilitator of excretion to a facilitator for import, and the previously externalised formate is then readmitted to the cell. This triggers the transcription of a number of genes involved in formate and pH homeostasis, including those encoding the enzyme formate hydrogenlyase (FHL-1) [8, 11].

FHL-1 is a membrane-bound enzyme made up of seven individual protein subunits [15]. It consists of a membrane arm, comprising two integral membrane proteins, and a peripheral (or catalytic) arm located at the cytoplasmic side of the membrane, comprising formate dehydrogenase-H (FDH-H) and hydrogenase-3 (Hyd-3). FDH-H is encoded by the *fdhF* gene and is a molybdenum- and selenium-dependent formate dehydrogenase [16]. Hyd-3 is a [NiFe] hydrogenase enzyme [17, 18] and is predicted to be the principal point of contact of the catalytic arm with the membrane arm [15, 18]. Under mixed-acid fermentation conditions, the two enzyme activities work together as a closed redox complex where FDH-H first oxidises formate to CO_2_ and then passes two electrons directly to the Hyd-3 active site where two protons are reduced to molecular hydrogen (H_2_), which can then diffuse away from the cell. Overall, therefore, FHL-1 carries out the disproportionation of formate to CO_2_ and H_2_ [11].

In this work, the physiology of H_2_ and formate metabolism in living intact cells has been revisited using powerful analytical spectroscopic techniques and isotopic labelling, which can monitor formation and consumption of compounds over time without invasive sampling of the batch culture [19, 20]. Surprisingly, under anaerobic fermentative conditions, isotopic labelling experiments revealed that exogenously-added formate behaved quite differently from endogenously-produced formate, being taken up and immediately disproportionated to CO_2_ and H_2_ while excretion of endogenous formate continued as normal. In addition, when *E. coli* was grown in our system in the presence of O_2_, it was possible to identify aspects of anaerobic metabolism (for instance formate and ethanol production) occurring at the same time as O_2_ uptake, indicative of the establishment of a micro-aerobic (micro-oxic) environment. Interestingly, under this micro-aerobic growth regimen, despite formate production and consumption being observed, no H_2_ production was detected unless exogenous formate was added to the culture, in which case H_2_ gas production proceeded in an FHL-1-dependent manner. This work demonstrates that FHL-1 can be assembled and remains active under microaerobic conditions, and gives confidence this cellular system could be further harnessed for biotechnological applications where O_2_ levels may fluctuate or be difficult to control.

## METHODS

### Bacterial strains and growth conditions

*E. coli* K-12 strains utilised in this study were MG1655 (F^−^, λ^−^, *ilvG*^−^, *rfb-50, rph-1* [1]) and MG16dZ (as MG1655 Δ*fdhF* [21]). Strains were transferred from −80 °C glycerol stocks, streaked on LB-agar plates and incubated overnight at 37 °C. Before each aerobic experiment, a 50 mL starter culture of LB medium was prepared in a sponge-capped tube, inoculated with a single colony and incubated for 16 h (37 °C, 200 rpm) to a typical OD_600_ of 1.2. For anaerobic experiments, a static, sealed tube filled almost to the top with medium to limit headspace, was prepared. Ultimately, 20 mL aliquots of the starter cultures were harvested by centrifugation and the cell pellets suspended in 20 mL of fresh M9 minimal medium ready for the experimental phase.

### Continuous spectroscopic monitoring of anaerobic fermentation

A combination of advanced non-invasive analytical spectroscopic techniques was used to monitor bacterial batch cultures *in situ* in a closed system [19, 20]. First, 230 mL of M9 medium was prepared in the round bottom flask with two side-arms. The flask was maintained at 37 °C using a temperature-controlled water bath under rapid stirring to enable efficient gas transfer. Next, the 20 mL anaerobic starter culture of *E. coli* prepared in M9 medium was added to the 230 mL of fresh M9 medium in the 2-neck round-bottom flask giving a typical starting OD_600_ of 0.1 and pH of 6.9. The culture was supplemented with 30 mM D-glucose (final concentration), and where appropriate, 20 mM deuterium (^2^H or D)-labelled formate (Sigma 373842, 99 % D) or 20 mM ^13^C-labelled formate (CK isotopes CLM-583, 99 % ^13^C) as potassium salts (final concentrations). The flask was then sealed and purged of O_2_ by alternating between evacuating the headspace and refilling with N_2_ at least five times.

While recording metabolic activity, the 2-neck flask containing the 250 mL *E. coli* suspension was held at 37 °C and rapidly stirred inside a thermostated water bath. From one neck of the flask, the 1425 mL headspace gas was cycled for gas-phase FTIR and Raman measurements by a peristaltic pump with a flow rate of 4.5 L/h. Production of ^12^CO_2_ and ^13^CO_2_, ethanol and acetaldehyde was quantified by gas-phase FTIR spectroscopy (Mattson Research Series, 0.4 cm^-1^ spectral resolution) with a home-built, 6 m pathlength multiple-pass White cell. Partial pressures were obtained by fitting experimental spectra to the sum of reference spectra from the HITRAN [22] and PNNL [23] databases. Gas-phase cavity-enhanced Raman spectroscopy (CERS) monitored O_2_, N_2_, ^12^CO_2_, ^13^CO_2_, and H_2_, HD and D_2_ isotopomers. In the CERS experiment, the 40 mW 636 nm cw-diode laser was enhanced by around four orders of magnitude in an optical cavity [24-26]. Scattered Raman light was analysed in a monochromator (Andor Shamrock SR163, DV420A-OE camera). Partial pressures were obtained from the Raman spectra after a calibration [27]. Using Henry’s law, pressures were converted into concentrations in solution. Using the ideal gas law, we estimate that 10 % CO_2_, 95 % acetaldehyde and 99.7 % ethanol were dissolved [20]. Under our conditions, less than 1% of dissolved CO_2_ was expected to be converted to carbonic acid and carbonates.

The liquid suspension was cycled from the second neck of the flask for *in situ* OD_600_ and liquid Raman measurements by a second peristaltic pump (4.5 L/h). The suspension was circulated through a sealed borosilicate tube for recording liquid-phase Raman spectra using a home-built spectrometer with a 532.2 nm, 20 mW laser (Lasos, GL3dT) and monochromator (Shamrock SR-750-A, DU420A-OE camera) [19, 20]. No interfering fluorescence was noticeable in M9 minimal growth medium. The water bending vibration at 1630 cm^-1^ was used to normalise decreasing Raman intensities as the turbidity of the bacterial suspension increased. A fit of the 820-1200 cm^-1^ region to reference spectra yielded the concentrations of glucose and the monobasic and dibasic phosphate anions of the phosphate buffer, and a fit of the 1300-1500 cm^-1^ region provided acetate and unlabelled and labelled formate concentrations [19, 20]. Although lactate has also some weak Raman peaks within our range, we have not been able to identify lactate metabolism products in this study above our detection limits, estimated to be in the low mM region (approximately 2 mM). Note that Raman scattering within the bacterial cells likely also contributes to the experimental spectra. However, as the sum volume of bacterial cells is much less than the volume of the liquid medium, it is assumed the experimental spectra correspond mainly to species in solution. The liquid suspension cycling loop contained also a 1 cm optical pathlength glass cuvette where the scattering of a red laser pointer was continuously monitored to obtain OD_600_ data after a calibration.

At the end of the bacterial growth period the cell suspension was harvested by centrifugation, washed and dried at 37 °C for 48 hours to record the dry biomass (typically around 200 mg). The endpoint pH of the spent culture medium was recorded using a Mettler Toledo SevenMulti pH meter to corroborate the spectroscopically calculated values.

### Continuous spectroscopic monitoring of aerobic respiration

A 20 mL aerobic starter culture, prepared in M9 medium, was added to 230 mL fresh M9 medium in a 2-neck round-bottom flask giving a typical starting OD_600_ of 0.1. Carbon sources were added as required and the flask was rapidly stirred at 37 °C. In this case, the headspace of the flask initially contained just air and was not purged with N_2_, and O_2_ levels in the closed system were monitored by CERS in addition to all other spectroscopic measurements.

### Calculation of changes in culture pH over time

The M9 medium phosphate buffer, composed of 47 mM HPO_4_^2-^ and 22 mM H_2_PO_4_^-^, absorbs H^+^ from excreted acids during bacterial growth by shifting HPO_4_^2-^ to H_2_PO_4_^-^. Liquid-phase Raman measurements were used to calculate the concentrations of phosphate anions, which in turn were used to calculate the pH *in situ* using a modified Henderson-Hasselbalch equation [19, 28].

### Data handling and visualisation

All experiments conducted in this study were repeated at least in triplicate, with all repeats showing essentially the same behaviour (see Supplementary Information). Figures in the main text are presented as single representative experiments. Partial pressures *p* (mbar) were converted to number of moles *n* by using the ideal gas law (*V* = 1.425 × 10^−3^ m^3^, *T* = 310 K) and correcting for the dissolved percentage calculated by Henry’s law. Note that *n* and *p* of CO_2_ displayed in the figures is scaled up to correct for the 10 % dissolved in solution.

## RESULTS

### Fermentation of glucose produces equimolar CO_2_ and H_2_

First, mixed-acid fermentation was studied using the well-characterised *E. coli* K-12 strain MG1655 [1] as a model system. An anaerobic starter culture of MG1655 was used to inoculate fresh M9 medium supplemented with 30 mM glucose (*i*.*e*., 7.5 mmol glucose in the 250 mL flask). Growth characteristics and fermentation products were quantified by CERS, FTIR and liquid Raman spectroscopy over 18 hours at 37 °C (Figure 1). Overall, these fermentation data can be divided in to at least three phases as defined by the behaviour of the endogenously-produced formate (Figure 1b). Following inoculation there is a typical lag phase lasting around 3 hours (Figure 1d). Then, as growth commences, net export of formate dominates until the 9 hour mark (Figure 1b, phase A), which coincides with the slowing of logarithmic growth (Figure 1d). During this first phase net formate accumulation peaks at 4 mmol in the extracellular medium (Figure 1b, phase A). Next, there is a decrease in formate as net import exceeds export (Figure 1b, phase B) and this continues until the 15 hour mark, by which time the cells have settled in to stationary phase (Figure 1d). Finally, formate is exhausted (Figure 1b, phase C).

**Fig. 1.**
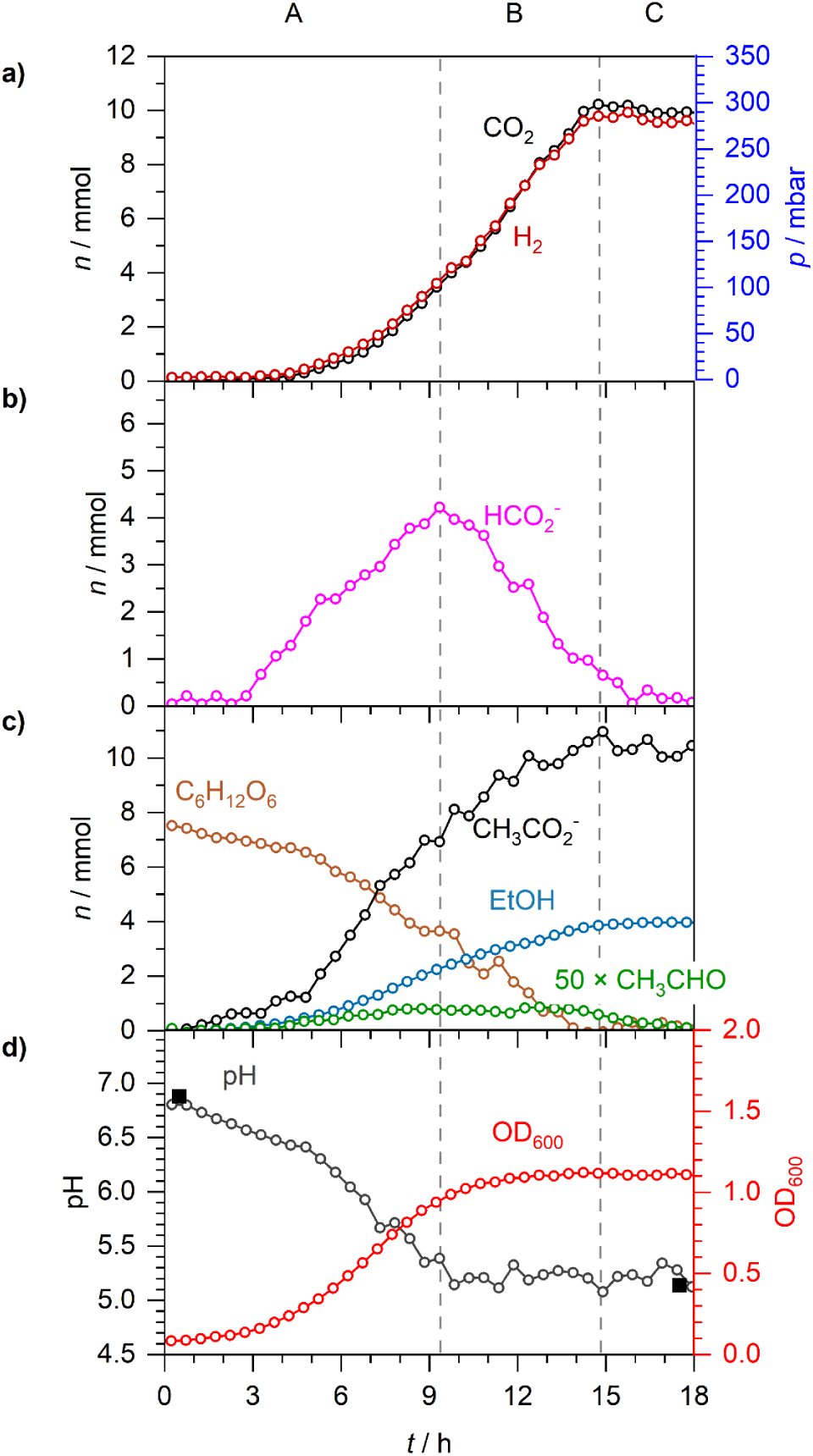
Anaerobic fermentation of glucose by *E. coli* K-12. Anaerobic fermentation by *E. coli* MG1655 during growth in M9 medium supplemented with 30 mM glucose. **A** to **C** denote three distinct phases: net formate excretion (A), net formate consumption (B) and formate depletion (C). **a)** Time-dependent number of moles (*n*) and equivalent partial pressures (*p*) of CO_2_ and H_2_. **b)** *n* of formate. **c)** *n* of acetaldehyde (×50), acetate, ethanol and glucose. **d)** Spectroscopically determined pH (open circles), externally measured pH (solid squares) and OD_600_.

Using this experimental approach, the activity of FHL-1 is evident throughout the growth phases until formate is finally exhausted at 15 hours (Figure 1a). Molecular hydrogen and CO_2_ are readily quantified in the gas phase and their production kinetics map precisely together, culminating in 10 mmol each of H_2_ and CO_2_ being produced (Figure 1a).

The liquid Raman spectroscopy allows the consumption of glucose to be followed, together with the accumulation of other fermentation products (Figure 1c). Acetate was produced to a final level of 11 mmol, and 4 mmol ethanol was generated (Figure 1c). Interestingly, trace amounts of acetaldehyde, the enzymatic intermediate between acetyl-CoA reduction to ethanol using NADH, was observed in the FTIR data and accumulated at a peak value of 18 μmol in phase B. From 14 to 18 hours, the remaining acetaldehyde was exhausted (Figure 1c).

### The fate of exogenous formate during fermentation

One powerful aspect of Raman spectroscopy is the ability to differentiate between isotopically labelled compounds. Formate is amenable to ^13^C- and D(^2^H)-isotope labelling studies due to the low natural abundances of carbon-13 (1.1 %) and deuterium (0.015 %). Figure 2 shows isotope shifts of the C-O stretching vibration for unlabelled formate (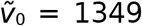 cm^-1^), formate-^13^C (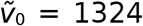 cm^-1^) and formate-D (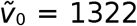 cm^-1^) in liquid Raman spectra. The isotope shift opens the door to fermentation experiments where exogenously-added ^13^C- or D-labelled formate could be readily distinguished from any endogenously-produced formate, which would remain unlabelled.

**Fig. 2.**
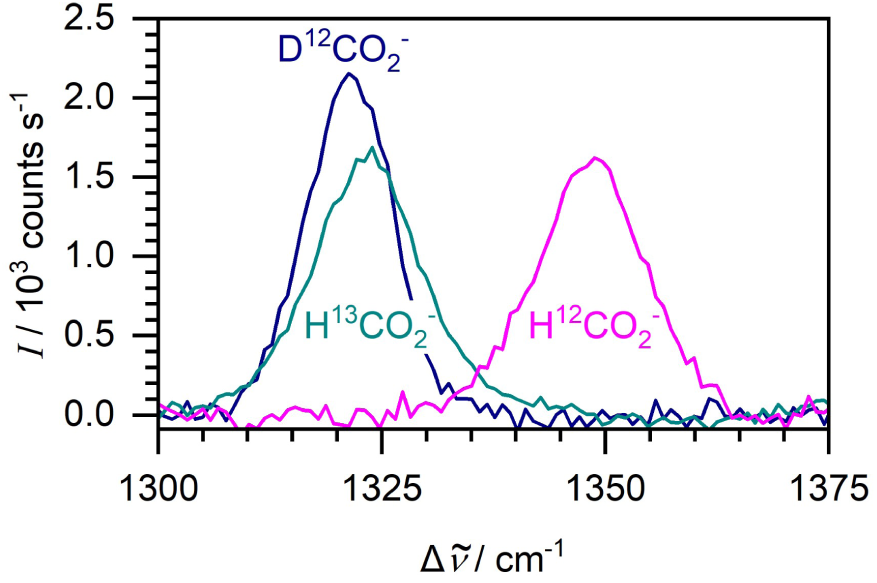
Raman spectroscopy can differentiate between isotopically-labelled formate anions. Liquid-phase Raman spectra of 100 mM unlabelled, D-labelled and ^13^C-labelled formate.

Arguably one of the most compelling aspects of formate metabolism is the membrane transport processes at play in this biological system. In order to examine the metabolism of formate added exogenously at the beginning of the growth phase, 20 mM (5 mmol in the flask) formate-D (as a potassium salt), together with 30 mM unlabelled glucose, was introduced into anaerobic batch cultures of *E. coli* MG1655 (Figure 3). Under these conditions, bacterial growth rates, and the amounts of acetate, ethanol and acetaldehyde produced, were comparable with the cells grown in the absence of exogenous formate (Figures 1,3). However, notable differences were observed in the formate, CO_2_, H_2_ and pH data (Figure 3).

**Fig. 3.**
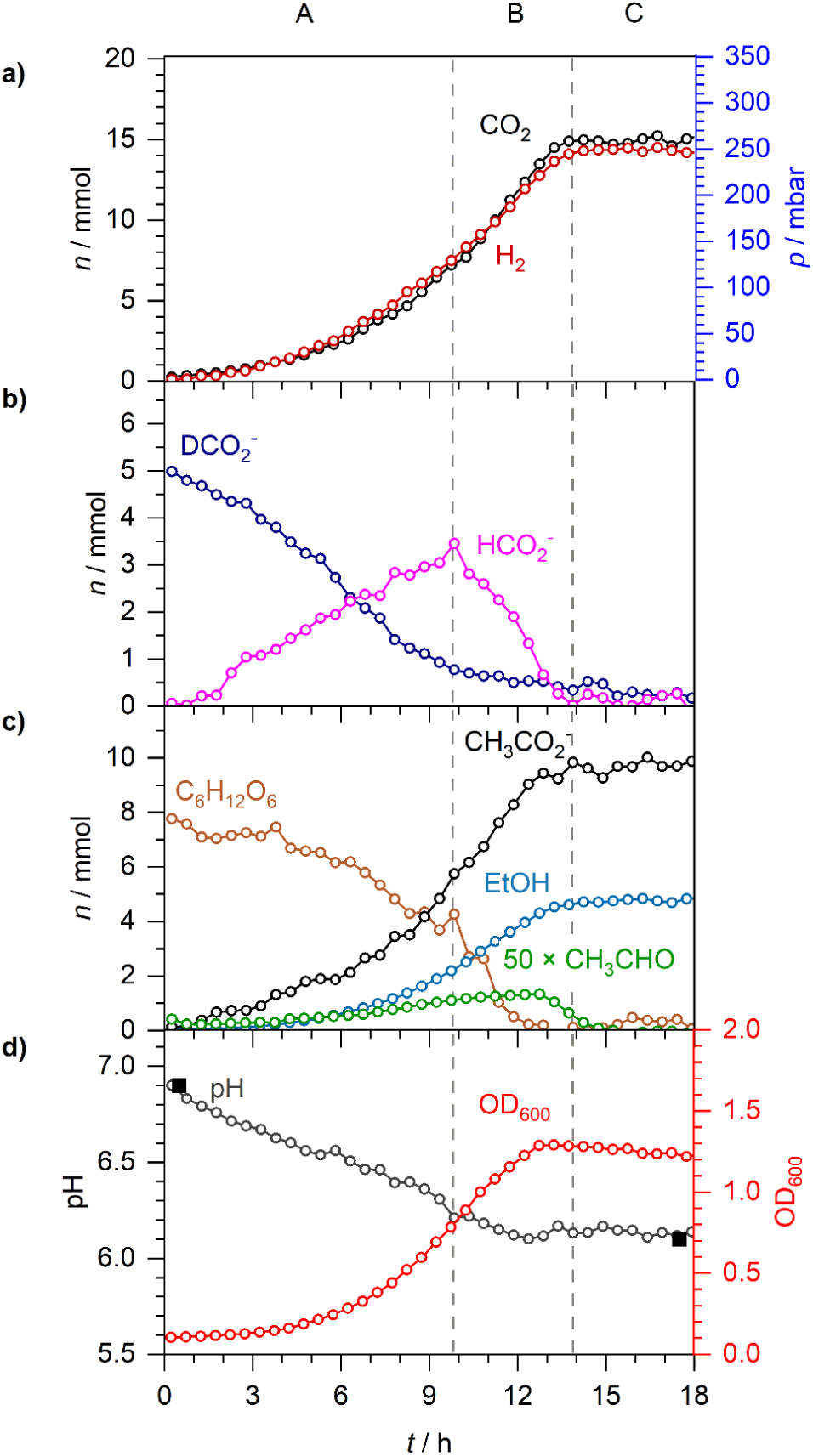
Exogenous formate-D is taken up by the cell, increasing CO_2_ and H_2_ production. Anaerobic fermentation by *E. coli* MG1655 during growth in M9 medium supplemented with 20 mM formate-D and 30 mM glucose. As before, **A** to **C** denote three distinct phases: net formate excretion (A), net formate consumption (B) and formate depletion (C). **a)** *n* and *p* of CO_2_ and H_2_. **b)** *n* of formate-D and formate. **c)** *n* of acetaldehyde (×50), acetate, ethanol and glucose. **d)** Spectroscopically determined pH (open circles), externally measured pH (solid squares) and OD_600_.

Metabolism of the external 5 mmol formate-D was observed to begin almost immediately (Figure 3b). Given the final amounts of H_2_ and CO_2_ produced in this experiment (15 mmol, Figure 3) compared to the amount derived from the experiment without exogenous formate (10 mmol, Figure 1), it is most likely that all of the external formate-D was disproportionated to CO_2_ and H_2_ by FHL-1. Note that although the oxidation of formate-D would produce CO_2_ and D^+^, no detectable levels of HD or D_2_ gas, which are also readily distinguished from H_2_ by CERS, were observed.

Remarkably, despite the presence of external formate-D, production, excretion and metabolism of endogenously produced unlabelled formate followed a familiar pattern (Figure 3b). In the initial phase, net export of formate was observed until around 10 hours and peaked at around 3.5 mmol (Figure 3b, phase A). There then followed a net decrease in formate concentration as import and disproportionation continued until the 14 hour mark (Figure 3b, phase B), by which time the FHL-1 reaction had ceased (Figure 3a) and the cells were in stationary phase (Figure 3d). Interestingly, at the 7 hour timepoint the extracellular concentrations of formate-D and unlabelled formate were identical (2.5 mmol), however the rates of excretion of endogenous formate and oxidation of formate-D continued largely unchanged (Figure 3b).

To explore further the relationship between exogenous and endogenous formate and the observed FHL-1 activity, a similar experiment using labelled formate-^13^C was performed (Figure 4). Note that ^13^CO_2_ can be distinguished from ^12^CO_2_ by CERS or FTIR. Here, 20 mM (5 mmol) formate-^13^C (as a potassium salt), together with 30 mM unlabelled glucose, was introduced into anaerobic batch cultures of *E. coli* MG1655 (Figure 4). The initial consumption of exogenous 5 mmol formate-^13^C mirrored the production of 5 mmol ^13^CO_2_, with the ^13^CO_2_ data perfectly overlapping with the formate-^13^C data if inverted (Figure 4). Production and excretion of endogenous unlabelled formate continued for 9-10 hours in this case (Figure 4b) and peaked at 4 mmol before net uptake and oxidation was observed (Figure 4b). This coincided with detectable levels of unlabelled ^12^CO_2_ being generated (Figure 4a). Indeed, the sum total of ^13^CO_2_ and ^12^CO_2_ overlapped almost exactly with the total H_2_ production levels, indicating that FHL-1 activity was responsible for the generation of these compounds.

**Fig. 4.**
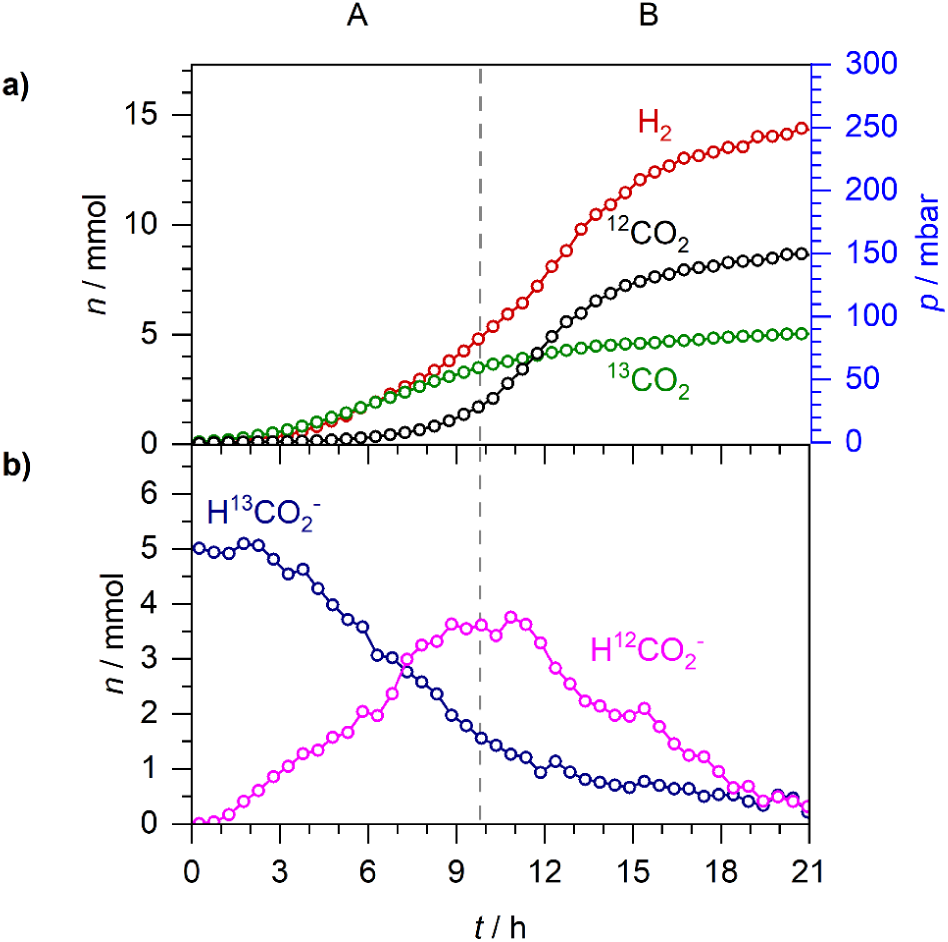
CO_2_ from exogenous and endogenous formate can be distinguished. Anaerobic fermentation by *E. coli* MG1655 during growth in M9 medium supplemented with 20 mM formate-^13^C and 30 mM glucose. **A** and **B** denote two distinct phases: net formate excretion (A) and net formate consumption (B). **a)** *n* and *p* of ^13^CO_2_, ^12^CO_2_ and H_2_. **b)** *n* of formate-^13^C and formate.

There were also notable differences in the final pH of the growth medium depending on the presence or absence of exogenous formate (Figures 1, 3). Addition of potassium formate salt did not acidify the growth medium (Figure 3d), hence the pH in both experiments started at 6.9 (Figures 1d, 3d). However, in the presence of exogenous formate-D the pH reduced from 6.9 to 6.1 over the course of the growth experiment (Figure 3d), compared to a drop in pH from 6.9 to 5.1 when the cells are grown with exogenous glucose only (Figure 1d).

### Metabolic behaviour under microaerobic respiratory growth conditions

We next set out to explore formate metabolism in aerobic cell cultures. First, an aerobic starter culture of *E. coli* MG1655 was prepared and used to inoculate 250 mL of M9 medium supplemented with 30 mM glucose. Note that this is a closed system that contains air in the vessel headspace but no way to replenish or regulate O_2_ content. Over the course of 24 hours, the cells were observed to grow, following an initial lag phase, to reach a peak OD_600_ value of 1.6 at around 9 hours (Figure 5d). Cell density was then observed to decline slightly over time, suggestive of cell death phase following the stationary phase (Figure 5d). Molecular oxygen (O_2_) was readily quantified in the headspace using our CERS system, and 8 mmol O_2_ was consumed in this closed system, with an equal amount of CO_2_ produced, over 24 hours (Figure 5a).

**Fig. 5.**
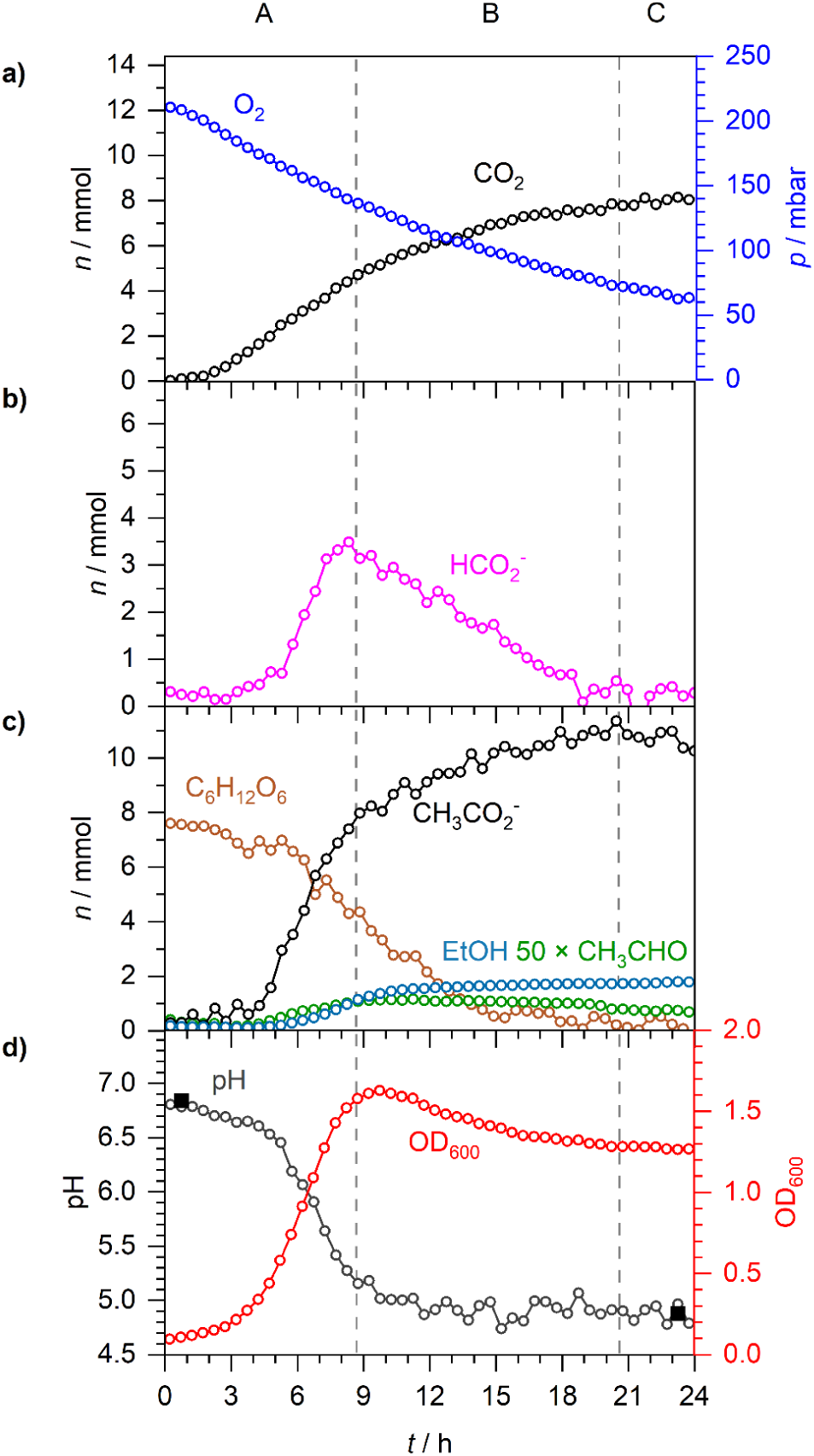
Evidence for microaerobic metabolism combining aspects of aerobic respiration with anaerobic fermentation. Aerobic respiration by *E. coli* MG1655 during growth in M9 medium supplemented with 30 mM glucose. **A** to **C** denote three distinct phases: net formate excretion (A), net formate consumption (B) and formate depletion (C). **a)** *n* and *p* of O_2_ and CO_2_. **b)** *n* of formate. **c)** *n* of acetaldehyde (×50), acetate, ethanol and glucose. **d)** Spectroscopically determined pH (open circles), externally measured pH (solid squares) and OD_600_.

Under these aerobic conditions, no detectable H_2_ production was observed at any point of the experiment (Figure 5a). However, there was evidence of anaerobic pathways occurring alongside aerobic respiration (Figure 5b, 5c), strongly suggesting a micro-aerobic environment was present. Indeed, endogenous formate metabolism was clearly evident in these data and Figure 5b could once again be readily divided in to phases A (0 - 8 hours), B (8 - 19 hours) and C (> 19 hours) based on this. Initial net formate production continued over 9 hours and peaked at around 3 mmol (Figure 5b, phase A). Next, the accumulated formate was consumed over the following 10 hours (Figure 5b, phase B), but no H_2_ was produced.

The excretion of acetic acid and formate caused the culture medium pH to decrease from 6.9 to 4.9 (Figure 5d, phase A), with the ultimate production of 11 mmol acetate and 2 mmol ethanol (Figure 5c). Taken altogether, it can be concluded that this experimental system is ideally suited to the study of micro-aerobic growth conditions, where aspects of anaerobic metabolism (especially formate and ethanol production) are functional alongside aerobic respiration.

### Addition of exogenous formate induces H_2_ production under aerobic conditions

Next, the micro-aerobic growth system was tested in the presence of excess exogenous formate. An aerobic culture of *E. coli* MG1655 was prepared in M9 medium containing 30 mM unlabelled glucose and 20 mM formate-D (Figure 6). Clear microaerobic physiology was observed, with formate, acetate and ethanol all being produced (Figure 6c), together with uptake of O_2_ from the gas phase (Figure 6a). As before, the observed endogenous formate metabolism could be divided in to three phases. First, endogenous formate is produced and excreted by the cell (across 0-9 hours), this time to a maximal level of 6 mmol (Figure 6b, phase A). Then the endogenously-produced unlabelled formate was oxidised or otherwise metabolised away (over 9-17 hours in this case) (Figure 6b, phase B), before a final phase was reached where formate was exhausted but other metabolic activities continued (Figure 6, phase C).

**Fig. 6.**
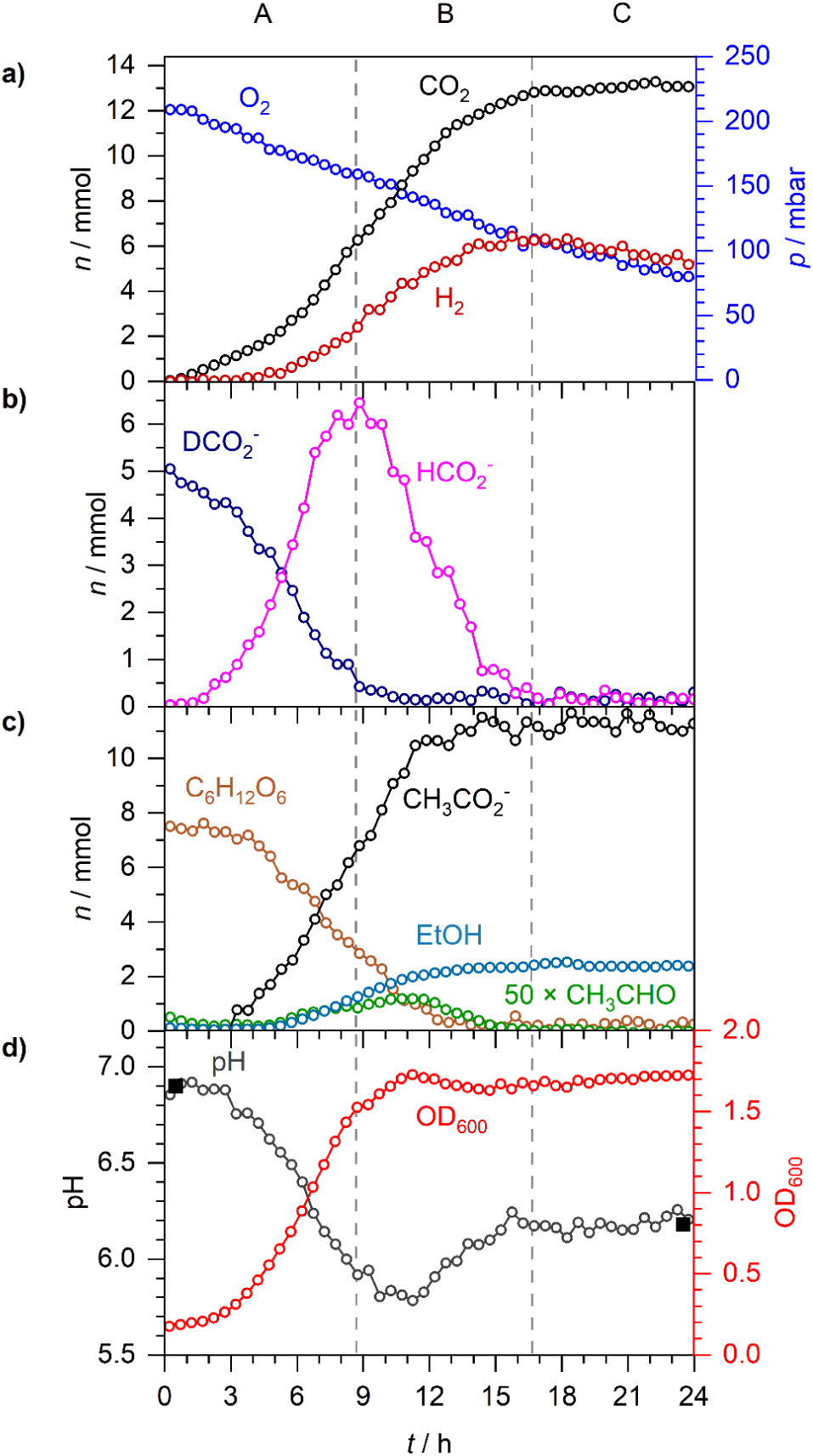
Exogenous formate induces H_2_ production during microaerobic respiration. Aerobic respiration by *E. coli* MG1655 during growth in M9 medium supplemented with 20 mM formate-D and 30 mM glucose. **A** to **C** denote three distinct phases: net formate excretion (A), net formate consumption (B) and formate depletion (C). **a)** *n* and *p* of O_2_, CO_2_ and H_2_. **b)** *n* of formate-D and formate. **c)** *n* of acetaldehyde (×50), acetate, ethanol and glucose. **d)** Spectroscopically determined pH (open circles), externally measured pH (solid squares) and OD_600_.

In this experiment, 5 mmol labelled formate-D was also added to the cultures from the outset. The exogenous labelled formate-D was consumed across phase A (Figure 6b). It is notable, however, that around the 5 hour timepoint the levels of formate-D and unlabelled formate appeared equal at 3 mmol each (Figure 6b). The kinetics of unlabelled formate production and formate-D uptake do not change at this point (Figure 6b). Most surprisingly, the addition of exogenous formate induced the production of H_2_ under aerobic conditions (Figure 6a). However, note that the molecular hydrogen was not observed in the data until around 5 hours in to growth (Figure 6a). Indeed, almost all of the 5 mmol of exogenous formate-D are consumed by the end of phase A, while only 2 mmol of H_2_ are produced in this time (Figure 6). An additional 4 mmol of H_2_ are produced in phase B, which also does not match with the consumption of at least 6 mmol of unlabelled formate in that time (Figure 6).

In order to test whether the aerobic H_2_ production observed in the presence of exogenous formate was as a result of FHL-1 activity a genetic approach was taken (Figure 7). The *E. coli* strain MG16dZ is a direct derivative of MG1655 carrying a complete deletion of the gene encoding the FDH-H component of FHL-1 [21]. An aerobic starter culture of *E. coli* MG16dZ was used to inoculate aerobic M9 medium supplemented with 20 mM formate-D and 30 mM glucose. Under these conditions, no detectable H_2_ production was observed at any point of the experiment (Figure 7). This strongly suggests that the H_2_ production induced by the addition of exogenous formate in the presence of O_2_ (Figure 6) was as a result of FHL-1 activity.

**Fig. 7.**
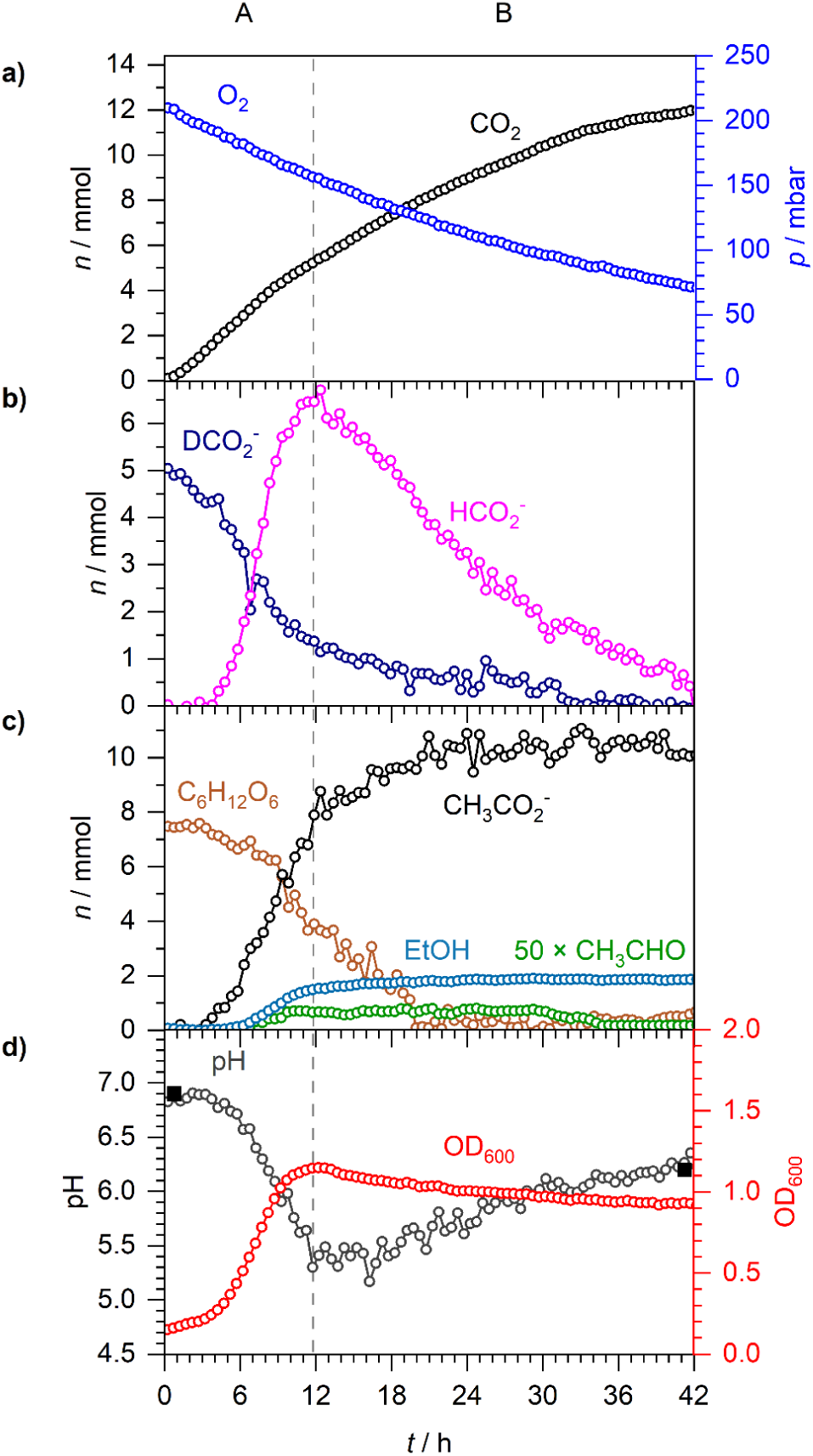
H_2_ production during microaerobic respiration is dependent upon FHL-1. Aerobic respiration by *E. coli* MG16dZ (Δ*fdhF*), which is FHL^-^, during growth in M9 medium supplemented with 20 mM formate-D and 30 mM glucose. **A** and **B** denote two distinct phases: net formate excretion (A) and net formate consumption (B). **a)** *n* and *p* of O_2_ and CO_2_. **b)** *n* of formate-D and formate. **c)** *n* of acetaldehyde (×50), acetate, ethanol and glucose. **d)** Spectroscopically determined pH (open circles), externally measured pH (solid squares) and OD_600_.

The microaerobic growth curve for MG16dZ (Δ*fdhF*) was similar to the parent strain MG1655, having a short lag before peaking at a OD_600_ of 1.1 at the 12 hour mark (Figure 7d). However, despite this the metabolic rates of the *fdhF* mutant are noticeably slower overall (Figure 7). For example, formate metabolism by MG16dZ (Δ*fdhF*) displayed only two clear phases even after 42 hours incubation (Figure 7). First, endogenous formate was produced from glucose across 0-12 hours, peaking at 6.5 mmol (Figure 7b, phase A). Then the endogenous, unlabelled formate was slowly consumed over the following 20 hours (Figure 7b, phase B). In addition, the majority (4 mmol) of the exogenous, labelled formate-D was consumed by 12 hours incubation (Figure 7b, phase A), by which time the cells had already concluded exponential growth (Figure 7d).

## DISCUSSION

In this work, the metabolism of *E. coli* K-12 in fermentative and microaerobic growth has been studied using in-line, non-invasive spectroscopic techniques. This allowed the consumption of glucose to be correlated with microbial growth and the production of numerous compounds, including H_2_ in the gas phase, to be quantified in real time. In addition, the ability of Raman spectroscopy to distinguish between isotopically-labelled compounds has allowed new insights in to formate metabolism to be made.

### Anaerobic mixed acid fermentation: evidence for efflux of formic acid

The physiology of *E. coli* growing under fermentative conditions is largely well understood [11], and one critical environmental factor driving gene expression and other physiological changes is the external pH. Using this experimental set-up it was possible to follow pH changes spectroscopically by monitoring the ratio of HPO_4_^2-^ and H_2_PO_4_^-^ [19, 28]. During exponential growth under fermentative conditions (Figure 1, phase A), 8 mmol acetate and 4 mmol formate (no lactate or succinate was able to be quantified in this experimental set-up within the approximately 2 mM detection limit, though they are expected to be present) accumulated in the external medium (Figure 1). Concomitantly, the external pH decreased from 6.9 to 5.1 (Figure 1), which can be explained if acetate and formate are excreted as acetic acid and formic acid (i.e. co-transport of the anions with protons). Indeed, the efflux of 12 mmol (48 mM) acidic protons (H^+^) in to the growth medium would be theoretically enough to overwhelm the 47 mM HPO_4_^2-^ component of the phosphate buffer used in these experiments, thus preventing further spectroscopic pH measurements if additional acidic products were excreted. Fortunately, the pH remains almost constant at 5.1 (Figure 1d), despite a further 3 mmol acetic acid being excreted after the peak of formate production at the end of exponential growth (Figure 1, phase B). This is because the reimport and disproportionation of 4 mmol formic acid essentially balances the efflux of acetic acid.

These observations are important when considering the structure and function of the FocA formate channel, and the physiology of formate metabolism and pH homeostasis. It is worth considering that formate (not formic acid) is the product of the pyruvate formate lyase (PFL) reaction at the cytoplasmic side of the membrane [29]. The PFL enzyme is anchored to, and possibly gating, the FocA channel in the cytoplasmic membrane such that it could be immediately excreted upon its synthesis [30, 31]. Efflux of formic acid (not simply formate) therefore requires co-transport of the anion with a new proton picked up from the cell cytoplasm. Such an event could not lead to charge separation across the membrane, but would result in the net movement of a proton from the cytoplasmic (*n*-side) to the periplasmic (*p*-side) of the membrane. Any possible bioenergetic advantages linked to this efflux are probably negated as *E. coli* attempts to maintain the pH of its cytoplasm. Under a range of external pH and growth conditions, the *E. coli* cell cytoplasm is maintained at 7.4-7.8 [32] thanks to a combination of ion transport events and compatible solute regulation [33]. Indeed, the FHL reaction itself seems to balance the pH of the growth medium (Figure 1) by the influx of formic acid (as opposed to formate alone) and its subsequent disproportionation to CO_2_ and H_2_. The influx of formic acid is further highlighted in our experiments with exogenously-added formate salts (Figure 3 and discussed further below).

### Separation of formic acid influx and efflux pathways during fermentation

The behaviour of formate, and its role in *E. coli* energy metabolism in general, has long been a source of intrigue in bacterial physiology. Endogenously-produced formic acid is initially excreted from the cell, presumably as *E. coli* has no real use for an excess of this C1 compound and it would perhaps be rapidly diluted, or metabolised away by other bacterial species, depending on the nature of the surrounding environment. Likewise, the presence of constitutively-expressed respiratory FDH-O originally posed a dilemma as endogenous formate is not made at all under fully aerobic conditions.

In this work, our experimental set-up clearly showed classic fermentative physiology where formic acid efflux was observed and the compound initially accumulated until the end of exponential growth, and was then metabolised away again (Figure 1b). In this case, net efflux of formic acid clearly continued as the external pH dropped from 6.9 to 5.5 (rather than stopping completely at a pH threshold), while H_2_ production was initially recorded at pH 6.5 (Figure 1a) and continued until formate was exhausted (Figure 1b).

In total, the anaerobic fermentation produced near equimolar quantities of 10 mmol CO_2_ and H_2_ (Figure 1a), 10 mmol acetate (Figure 1c), and 4 mmol ethanol from 7.5 mmol glucose in the initial medium. It is likely that succinate and lactate were also produced, but below the detection limits of our apparatus (*ca*. 2 mM). In theory, 7.5 mmol glucose could generate 15 mmol pyruvate from glycolysis. In turn, this could lead to 15 mmol acetyl-CoA (and 15 mmol formate) that could either be further reduced to ethanol or used for ATP production with an end product of acetate. The combined totals of acetate and ethanol are close to the theoretical (Figure 1), however 15 mmol of formate would be expected to generate 15 mmol of H_2_ and CO_2_ *via* FHL-1, while we recorded only 10 mmol for each (Figure 1a). It is possible that some formate may be retained within the cells (either periplasm, cytoplasm, or both), but at a level that cannot be efficiently oxidised by FDH-H (the *K*_m_ for formate for this enzyme is 10 mM [8]) or that it is being used for other biochemical pathways in the growing culture [34].

The ability of Raman spectroscopy to distinguish between isotopically-labelled compounds allowed some new observations of formate and H_2_ metabolism. First, it is worth documenting that the addition of formate-D to the culture did not lead to HD or D_2_ production. This is not altogether surprising as there would need to be an unusual mechanism of linking the FDH-H and Hyd-3 reactions within FHL-1 for this to be the case. Otherwise, the released D^+^ is too dilute at 20 mM (5 mmol) compared to the concentration of the bulk H^+^ in the aqueous phase (the concentration of pure H_2_O is 55.5 M). Second, under these anaerobic conditions all of the extra formate-D was disproportionated to H_2_ and CO_2_. This is evidenced by the fact that 15 mmol of both H_2_ and CO_2_ was recorded upon the addition of an extra 5 mmol of formate-D (Figure 3a), compared with 10 mmol each for the experiment with no exogenous formate (Figure 1a). Further corroboration was forthcoming in Figure 4a, where addition of 5 mmol of formate-^13^C to the culture led to 5 mmol ^13^CO_2_ being produced and an extra 5 mmol of H_2_ overall. Moreover, a kinetic analysis of the ^13^CO_2_ and ^12^CO_2_ data shows that the rate of CO_2_ production is consistent with being linearly dependent on the formate concentration and *E. coli* abundance (estimated by the OD_600_), without any apparent preference for formate-^13^C or unlabelled formate. As formate-^13^C was more abundant than endogenous formate in the extracellular medium to begin with, ^13^CO_2_ production was observed before ^12^CO_2_ production. Then, as formate-^13^C was consumed and endogenous formate was excreted, the rate of ^13^CO_2_ production decreased while the rate for ^12^CO_2_ increased (Figure 4a). This is possibly best viewed at the 7 hour timepoint when there were equal amounts (3 mmol) formate-^13^C and unlabelled formate in the extracellular medium (Figure 4b), and ^12^CO_2_ and ^13^CO_2_ have similar rates of production (Figure 4a).

The behaviours of the exogenously-added and endogenously-generated formate deserve some further discussion. Using isotopically-labelled formate, we have been able to observe both influx and efflux of formic acid occurring at the same time (Figures 3, 4). Indeed, the externally-added labelled formate and internally-produced unlabelled formate appear to behave as different substrates in this system, which could either be a subtle kinetic isotope effect or could be evidence for different molecular mechanisms of uptake (influx) and export (efflux) of formate. For instance, export of endogenously-produced unlabelled formate displays essentially the same behaviour, with similar rates and maximal levels observed, regardless of the presence or absence of external formate-D (Figures 1, 3). The maximum rate of formate efflux (normalised by the OD_600_), in the absence of externally-added formate, was calculated at 34 ± 3 μmol OD^-1^ min^-1^. When 5 mmol formate-D was added to the growth medium from the outset, the maximal rate of unlabelled formate efflux was calculated at 36 ± 7 μmol OD^-1^ min^-1^. For comparison, the maximum rate of D-formate influx was determined 111 ± 12 μmol OD^-1^ min^-1^, and the rate of re-import of unlabelled formate in stationary phase was 13 ± 5 μmol OD^-1^ min^-1^. Indeed, the relatively high rates of external formate influx during exponential growth are remarkable. It is worth considering that influx of formic acid is intimately linked with FHL activity [35], and that FHL activity is intimately linked, by an unknown mechanism, to the ATP levels in the cell [36]. It seems clear, given these data, that glucose-replete fast-growing cells contain greater physiological FHL activity than cells in the stationary phase.

Taken altogether, this gives extra weight to the hypothesis that PFL (which generates the internal formate from pyruvate) is physically attached to the N-terminus of the FocA formate channel and is gating it in ‘efflux mode’ such that formate production and formic acid efflux are intimately linked [30]. Such a model would effectively maintain the cytoplasmic formate concentration at close to zero with the immediate efflux of the product linked to its synthesis. However, in our experimental system, we already have formate-D at 5 mmol (above the maximal level of 4 mmol that the ‘natural’ system can achieve under these conditions) outside the cell. The influx of exogenous formate-D is coupled directly to its disproportionation to CO_2_ and H_2_ by FHL-1 and this appears to occur in parallel to efflux.

It is worth considering how efflux and influx of formic acid could occur at the same time in such a cellular system. A large body of work points to the FocA membrane protein as being primarily responsible for both efflux and influx of formic acid [11]. It seems unlikely that a given individual protein could both export and import formic acid simultaneously. Instead, it should be considered that a certain number of FocA channels in the cell could be gated for efflux only, for example those in physical contact with PFL, while a second pool of FocA channels could be gated for influx only. This perhaps also implies different routes of formate or proton translocation within the FocA protein under the different gated states. Indeed, site-directed mutagenesis of *focA* has identified amino acid residues that are important for formic acid influx, but not efflux. For example, substitution of *E. coli* FocA His-209 by either Asn or Gln locked the channel in efflux mode only [37]. This hypothesis would also stand if the monoculture of *E. coli* actually contained two distinct populations of cells, one carrying out formic acid efflux and another performing the FHL reaction linked to influx. Such bimodal phenotypic heterogeneity is common in bacterial populations [38].

In both experiments where exogenous formate was added (Figures 3, 4), formic acid consumption and H_2_/CO_2_ production began almost immediately with very little lag time. This was most likely because the starter cultures were prepared anaerobically and the cells would have a certain level of FHL-1 already present. It is notable, however, that external medium pH does not become as acidic when spiked with exogenous formate compared with normal glucose fermentation conditions – in the former the pH decreases from 6.9 to 6.1 (Figure 3) and in the latter the pH drops from 6.9 to 5.1 (Figure 1). This could be giving additional evidence as to how the FocA formate channel is working. A compelling model for FocA function is as an anion-proton symport system, where formic acid (HCOOH) would be either excreted or re-imported in to the cell [39-41]. In such a model the efflux of formic acid would acidify the medium (as was seen here, Figure 1), but conversely the subsequent influx of formic acid would de-acidify the medium and there may be clear evidence for that in Figure 3. Here, we added exogenous formate-D to the medium as a potassium salt (Figure 3). The potassium formate-D is not an acid and did not affect the starting pH of the experiment, which remained steady at 6.9 (Figure 3). Under the anion-proton symport model, influx of the 5 mmol extra, external formate-D would also bring with it 5 mmol of H^+^ across the membrane in to the cell from the bulk external medium, thus increasing the pH of the growth medium and helping to balance the pH of the external medium against the continued efflux of acetic acid and unlabelled formic acid.

### Microaerobic respiration

Following the fermentation experiments, attention then turned to the metabolism of formate under aerobic conditions. First, an *E. coli* starter culture already performing aerobic respiration was prepared and the experimental system was assembled with air in the culture headspace (200 mbar partial pressure is ∼20% O_2_, Figure 5). Over the course of the experiment, 8 mmol of O_2_ was consumed and 8 mmol of CO_2_ was produced – an apparent respiratory quotient of 1.0 indicative of aerobic respiration of glucose. However, clear aspects of anaerobic metabolism were also evident, including acetate and ethanol production, as well as the efflux and influx of endogenous formic acid (Figure 5b). Indeed, formate efflux/influx behaviour was almost identical to that observed for the fermenting cells (Figure 1) where 10 mmol each of H_2_ and CO_2_ were also generated. In the presence of O_2_, however, no H_2_ was observed whatsoever (Figure 5). In this case the formate was likely ultimately oxidised by the respiratory enzyme FDH-O in the periplasm [10], thus generating the 8 mmol of CO_2_ observed. Moreover, if all of the CO_2_ production is attributable to formate respiration, then it follows that half of the O_2_ consumption is attributable to formate oxidation in the periplasm. Taken altogether, in this experimental set-up we conclude the culture was clearly performing microaerobic respiration (Figure 5).

### FHL-1 activity is inducible under microaerobic conditions

Having established a micro-aerobic growth environment, the final experiment involved the study of the physiology of external formate metabolism under these conditions. Somewhat surprisingly, the addition of exogenous formate-D to the culture at the outset led to the production of H_2_ in an FHL-1-dependent manner (Figures 6, 7). Unlike the anaerobic fermentation experiments (Figure 3), however, there was a notable 5 hour delay in the onset of H_2_ production (Figure 6). This is most likely because, using an aerobic starter culture, the cells had to synthesise FHL-1 *de novo* in response to the external formate-D. Indeed, the formate-D was almost completely consumed by aerobic respiration before the H_2_ production began (Figure 6). However, sufficient endogenous, unlabelled formate was produced that enabled the ultimate production of 6 mmol of H_2_ and an extra 6 mmol of CO_2_ *via* FHL-1 (Figures 6, 7). Indeed, the ability of FHL-1 to function in the presence of 10-20% O_2_ in the headspace is quite remarkable, but, together with the obvious presence of numerous other O_2_-sensitive enzymes including PFL, the data is probably telling us that the O_2_ tension in the cell cytoplasm is very much lower.

It is pertinent to note here that there is continued consumption of O_2_ in stationary phase that is at roughly the same rate as consumption of the endogenous H_2_ (Figure 6a, phase C). Conversely, under strict anaerobic conditions, the H_2_ produced by FHL-1 remains largely steady (Figures 1, 3). It is tempting to speculate that at least some of this activity represents an example of ‘hydrogen cycling’ where H_2_ gas produced by FHL-1 is re-oxidised by the respiratory hydrogenases (Hyd-1 and/or Hyd-2) linked directly to the reduction of O_2_ to water [18]. Note, however, that continued O_2_ consumption up to 42 hours was observed in the absence of H_2_ (Figure 7), suggesting that other endogenous electron donors continue to be functional long after glucose is exhausted.

The H_2_ production induced by exogenous formate under microaerobic conditions is undoubtedly due to FHL-1 (Figure 7). However, the isogenic *fdhF* mutant has a phenotype that exhibits surprising weaknesses compared to the parent strain. Oxidation of both exogenous and endogenous formate is very slow – for example, for external formate-D the MG1655 parent strain consumed 4 mmol over 9 hours (Figure 3b), while for the FHL-1 mutant this process lasted 12 hours (Figure 7b). Moreover, the growth medium of the mutant strain retained significant amounts (1 mmol) of endogenously-produced formate even after 42 hours of incubation (Figure 7b).

The behaviour of formate and O_2_ respiration was also different between the mutant and the parent strain. While glucose was abundant from 0 - 18 hours, 8 mmol CO_2_ was produced and O_2_ decreased by 4 mmol, suggesting that all of the O_2_ respiration observed is linked to formate oxidation (Figure 7a). However, from 18 to 42 hours the glucose was depleted and endogenous formate could no longer be produced, hence the 4 mmol formate remaining was slowly oxidised to increase CO_2_ from 8 to 12 mmol, with O_2_ decreasing from 8 to 4 mmol at the same time, suggesting additional endogenous electron donors were coming into play in stationary phase.

### Concluding remarks

In this work, the formate and H_2_ physiology of *E. coli* has been examined by powerful, non-invasive Raman and FTIR spectroscopy. This project has produced further evidence that FHL-1 is part of a formic acid detoxification system involved in pH homeostasis. The data support an anion-protein symport model for formate transport across the cell membrane, and provide evidence for at least two separate routes, or mechanisms, for formic acid influx and efflux. Finally, we provide evidence that FHL-1 can be assembled and functional under microaerobic conditions, which could remove some barriers to biotechnological applications.

## SUPPLEMENTARY MATERIAL

Additional figures with all replicates performed are provided in the Supplementary Material.

## AUTHOR STATEMENTS

GDM was a Postgraduate Research Student who designed all experiments, conducted the research, analysed data, prepared figures for publication, and wrote the paper. FS analysed data and wrote the paper. MH conceived the project, assembled the research team, designed the research, conducted the research, supervised the research, analysed data, and wrote the paper.

## Supplementary Material

### S.1. Repeats of Fig. 1

**Fig. S1.**
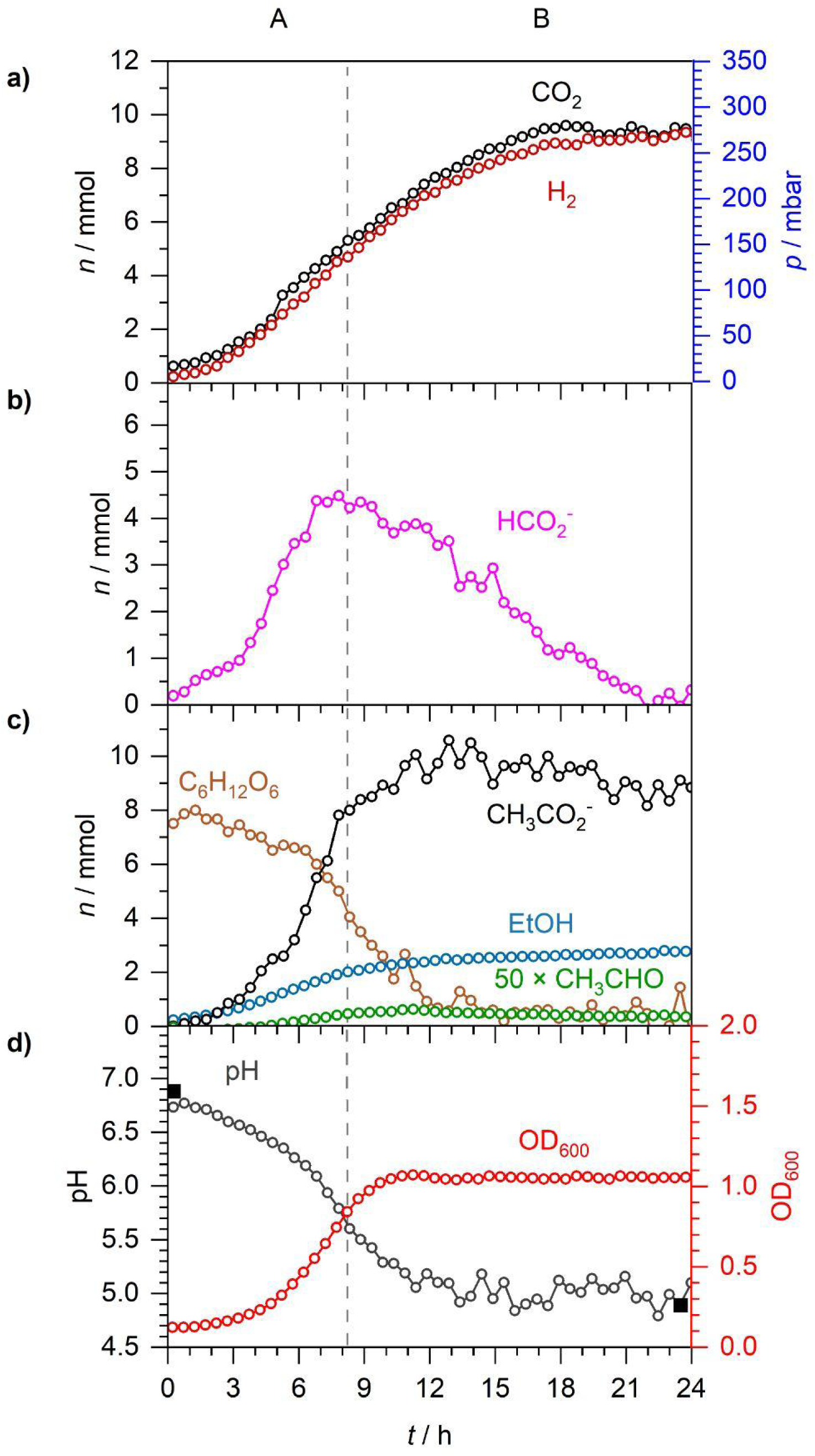
A repeat of Fig. 1.

**Fig. S2.**
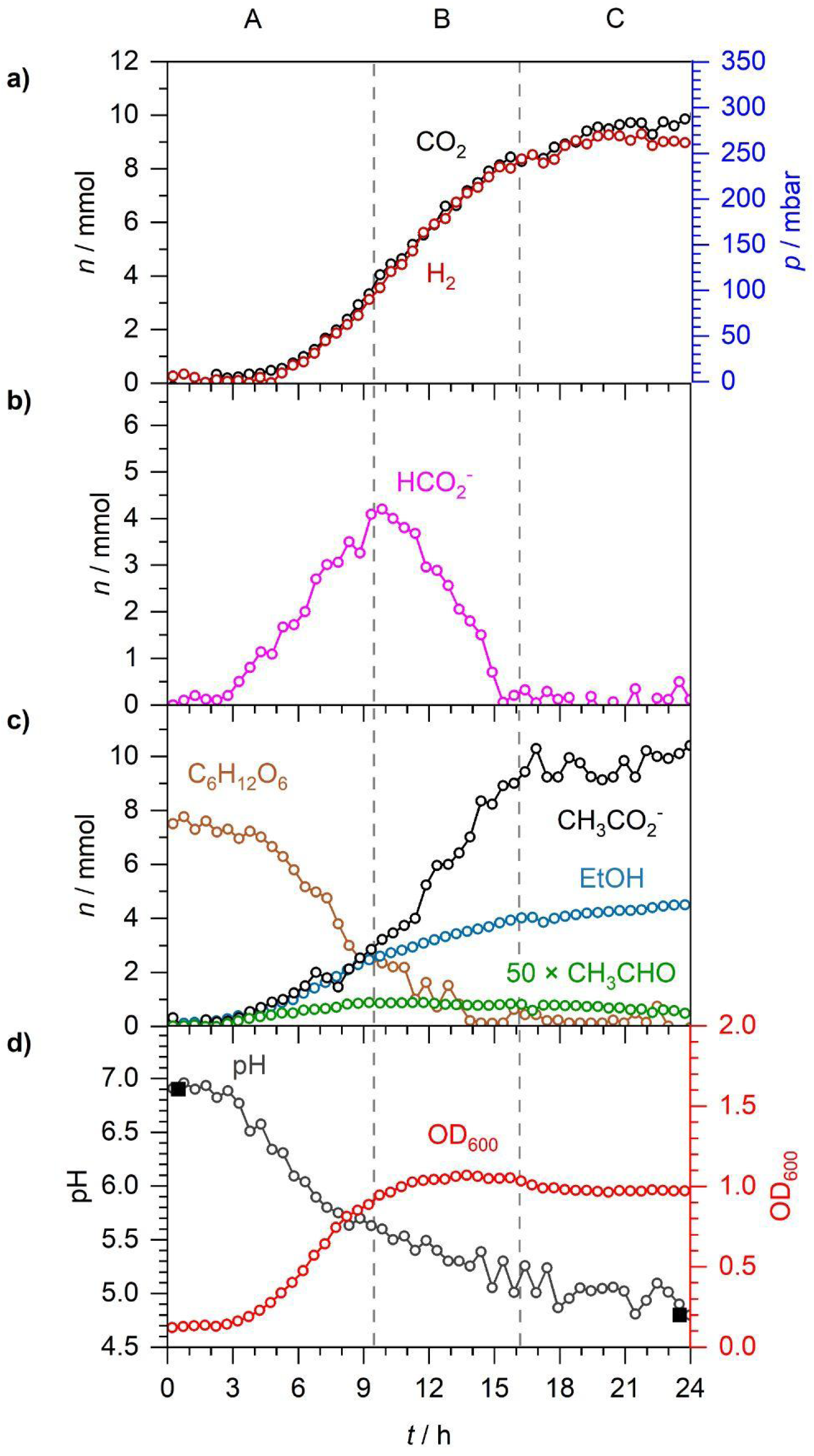
A repeat of Fig. 1.

### S.2. Repeats of Figs 3 and 4

**Fig. S3.**
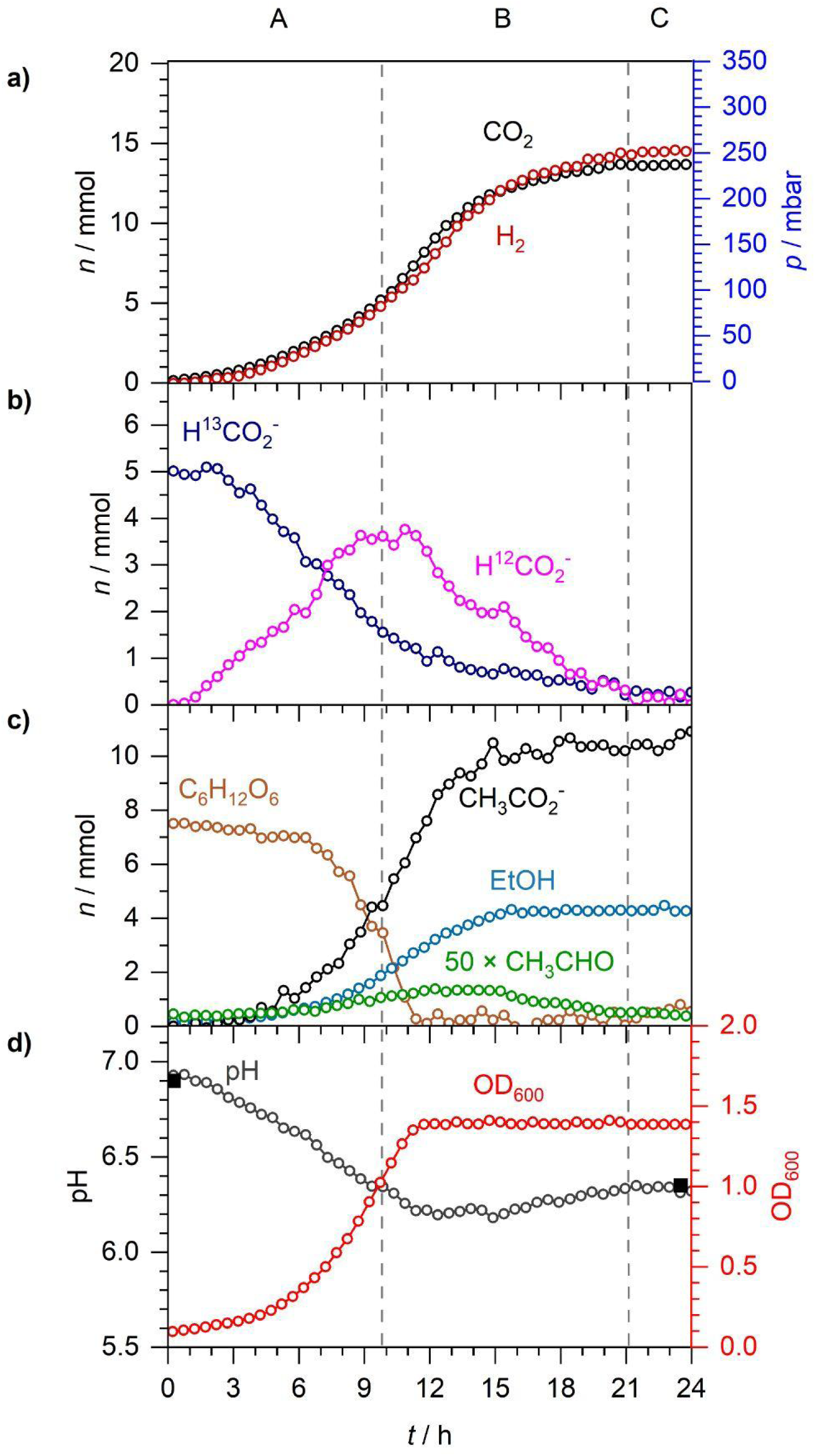
The same experiment as in Fig. 4 but with all measured data displayed. Here, CO_2_ is the sum total of ^13^CO_2_ and ^12^CO_2_. This also serves as a repeat of Fig 3.

**Fig. S4.**
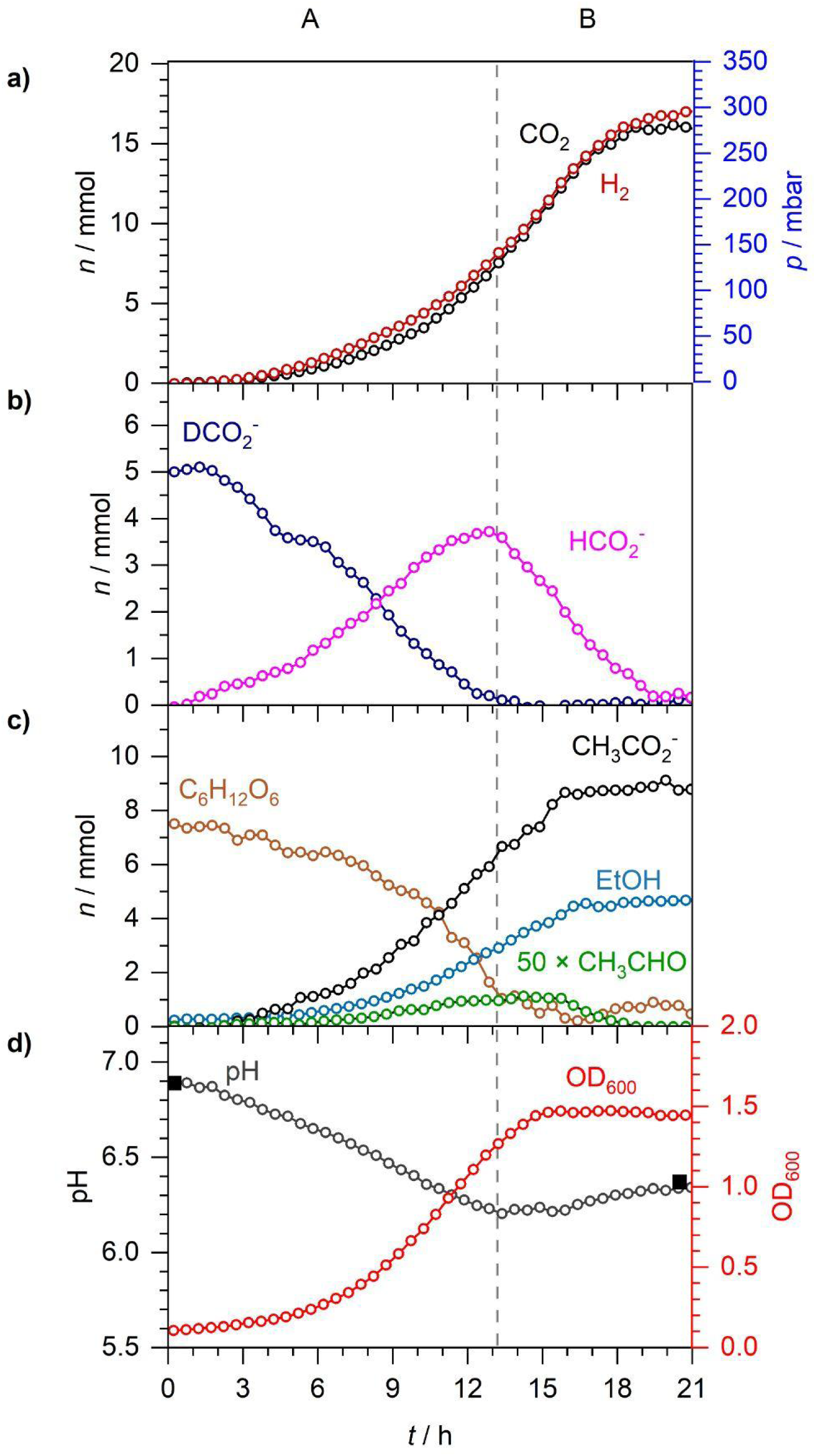
A repeat of Figs 3 and 4.

### S.3. Repeats of Fig. 5

**Fig. S5.**
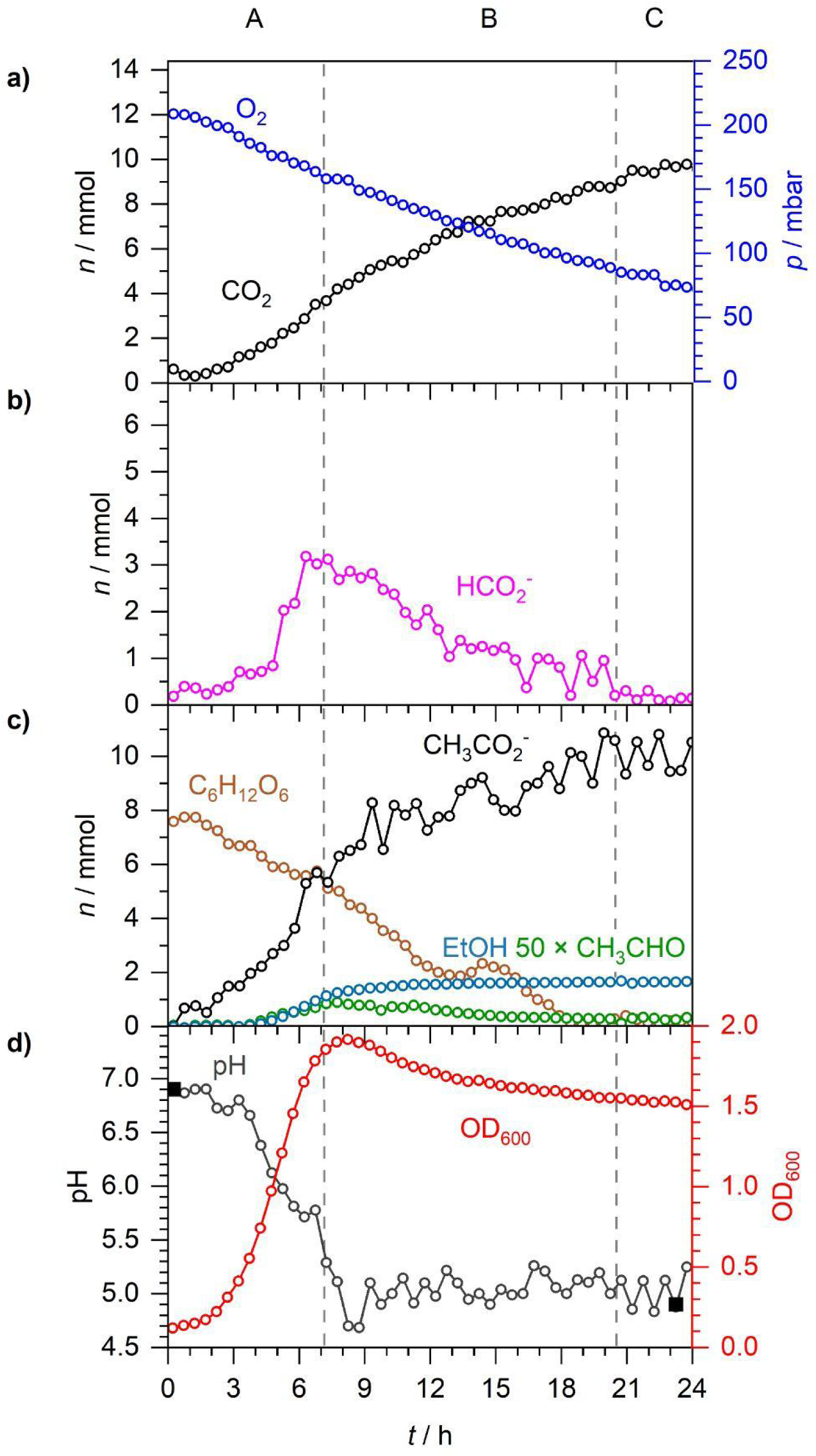
A repeat of Fig. 5.

**Fig. S6.**
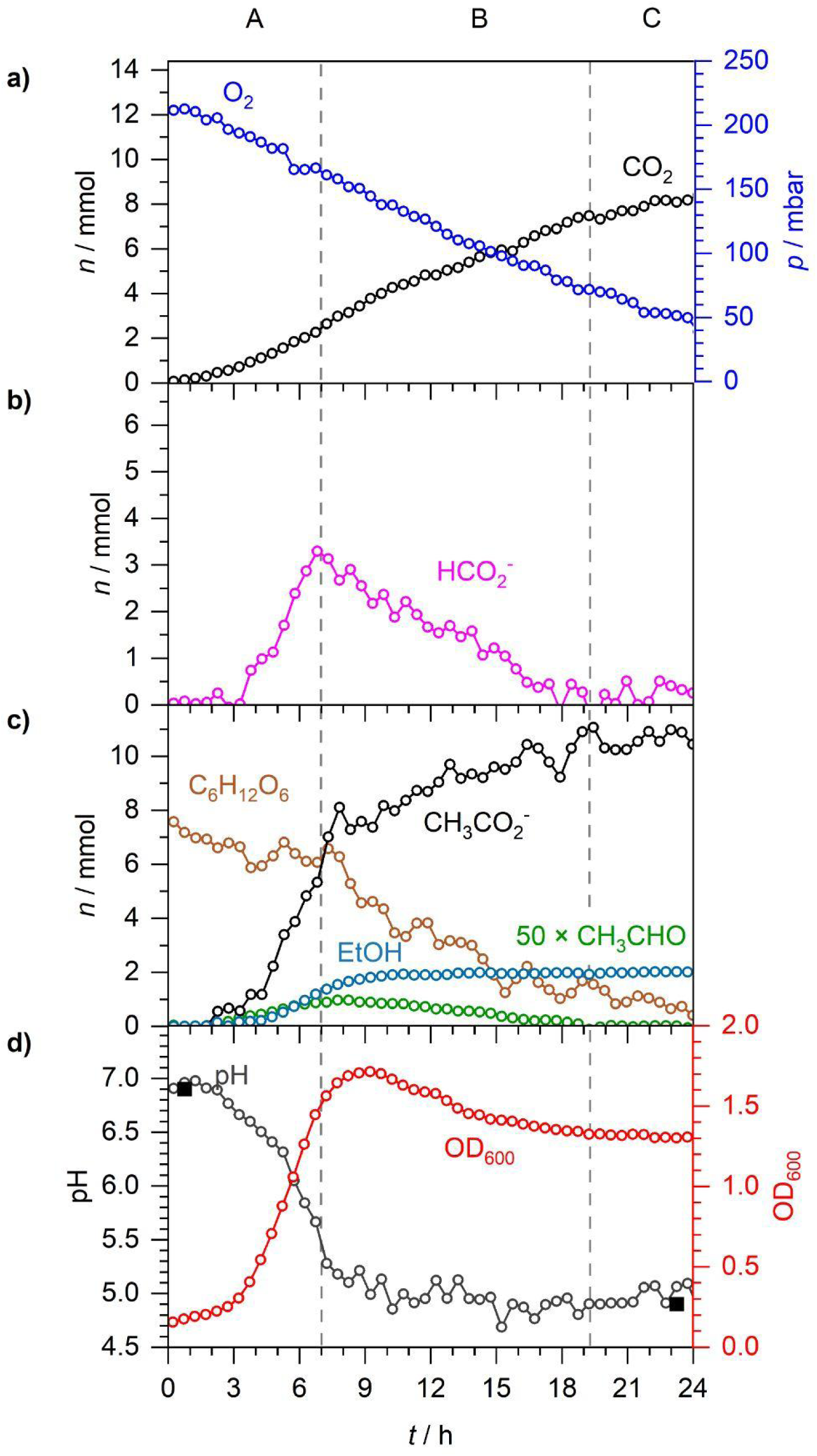
A repeat of Fig. 5.

### S.4. Repeats of Fig. 6

**Fig. S7.**
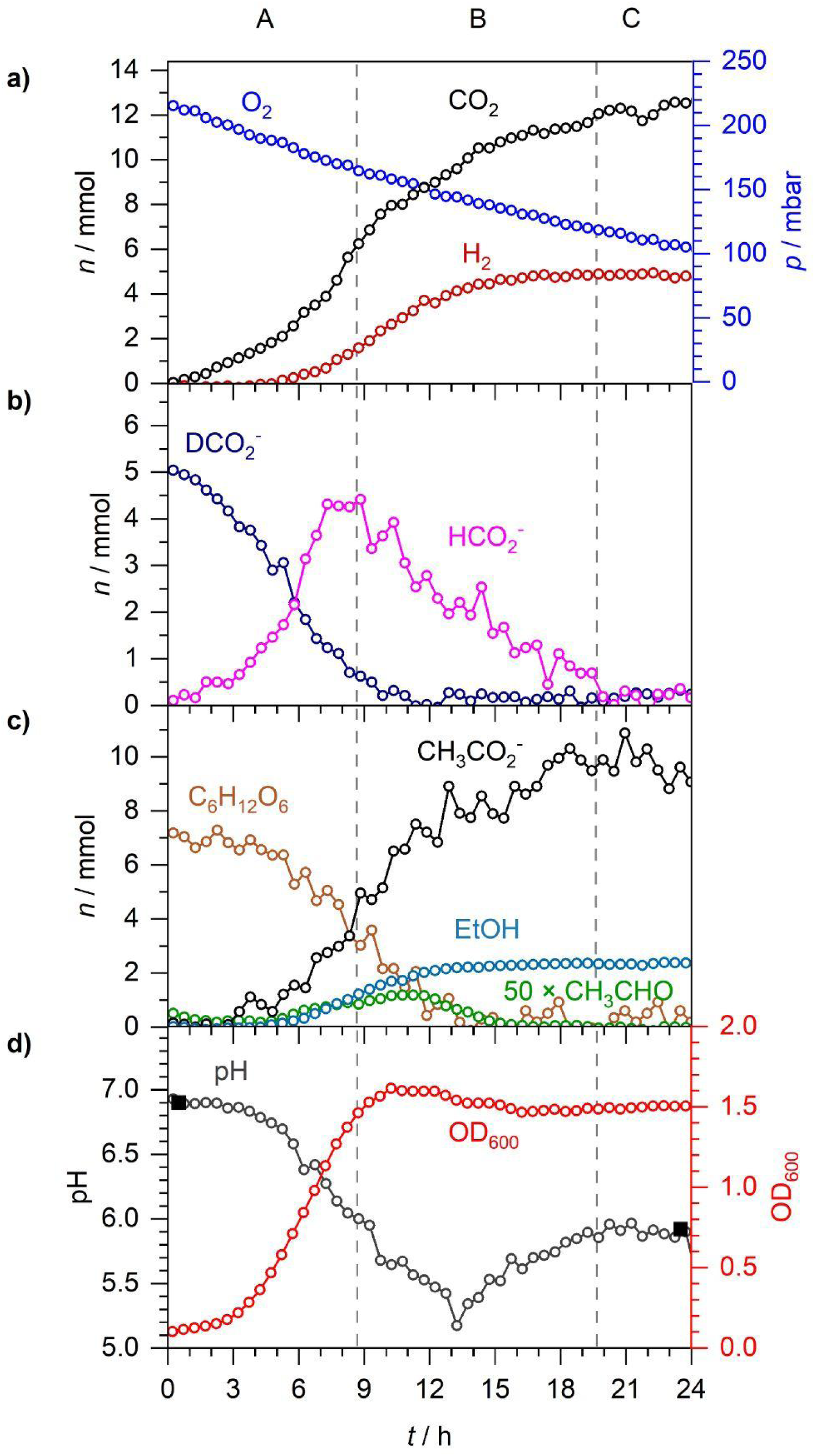
A repeat of Fig. 6.

**Fig. S8.**
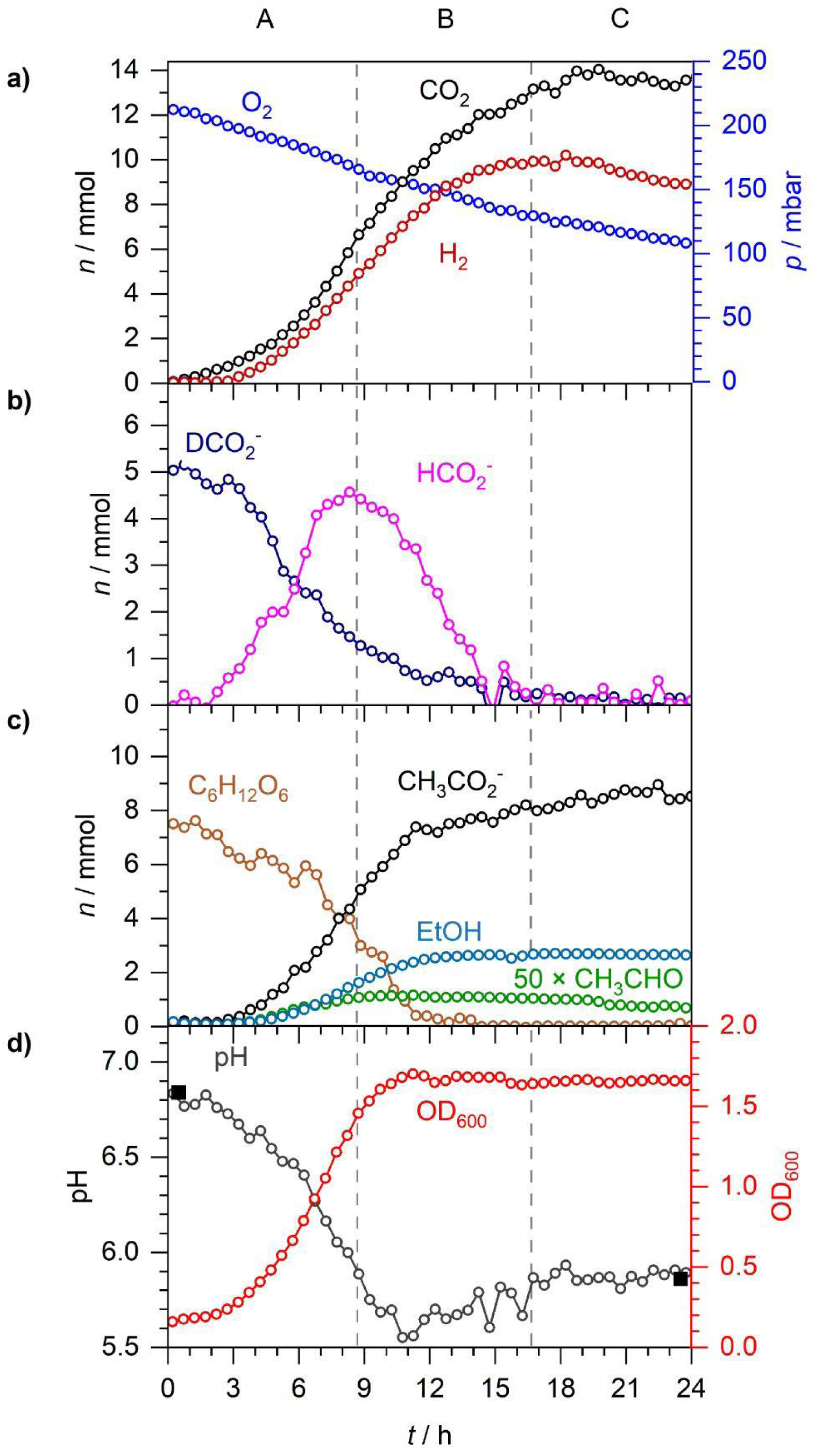
A repeat of Fig. 6.

### S.5. Repeats of Fig. 7

**Fig. S9.**
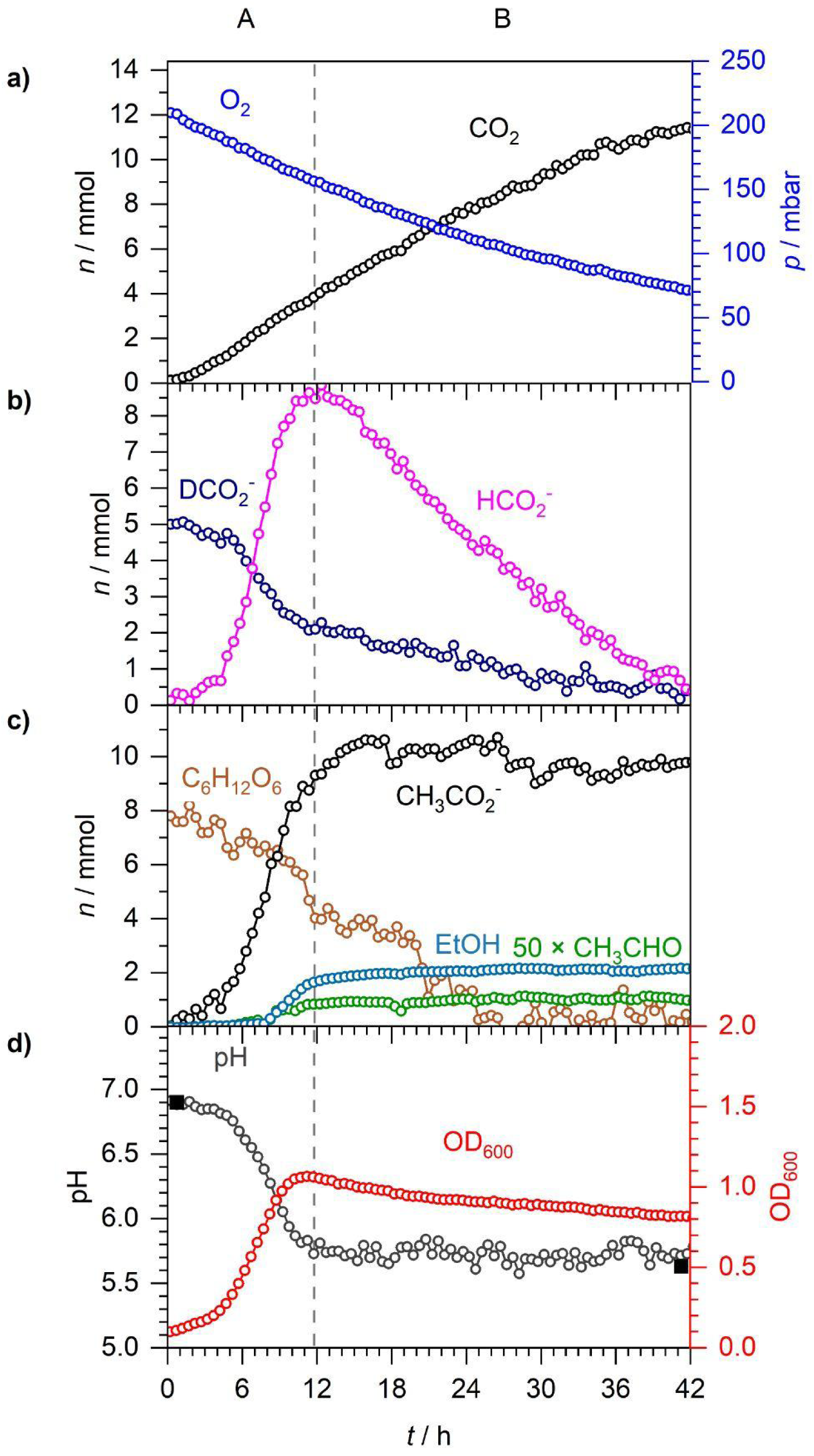
A repeat of Fig. 7.

**Fig. S10.**
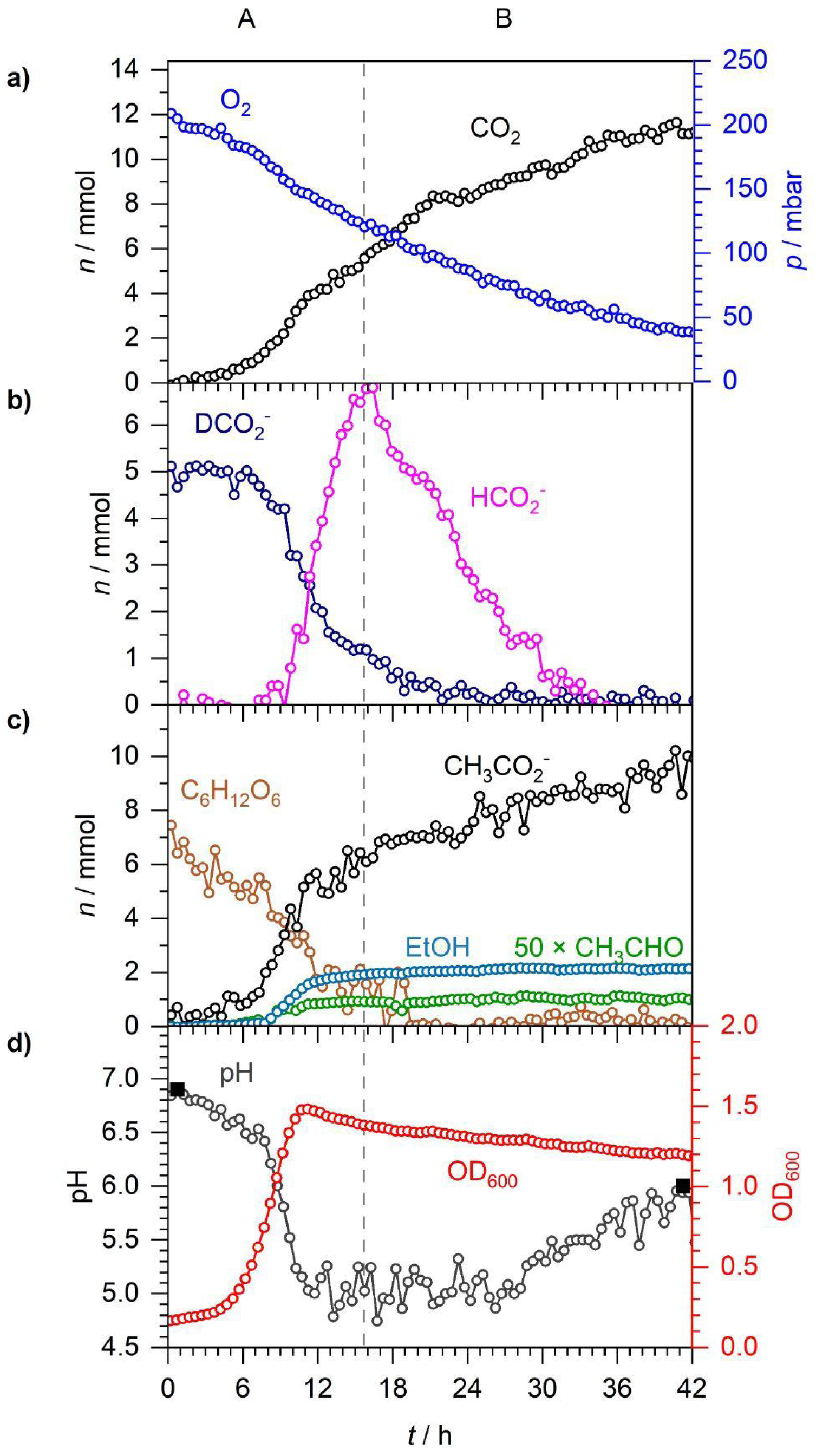
A repeat of Fig. 7.

